# TRANSCRIPTOMIC ANALYSIS OF THE RELATIONSHIP BETWEEN PANCREATIC DUCTAL ADENOCARCINOMA STROMAL SUBTYPES AND NEURAL INVASION

**DOI:** 10.1101/2025.01.13.632859

**Authors:** Saliha E. Yildizhan, Ihsan E. Demir, Elif Arik Sever, Didem Karakas Zeybek, O. Ugur Sezerman, Güralp O. Ceyhan

**Affiliations:** Department of Biostatistics and Medical Informatics, School of Medicine, Acibadem Mehmet Ali Aydinlar University, Istanbul, Turkey; Department of General Surgery, HPB-Unit, Acibadem Mehmet Ali Aydinlar University School of Medicine, Istanbul, Turkey; Department of Surgery, Klinikum rechts der Isar, Technical University of Munich, School of Medicine, Munich, Germany; 4SFB 1321, Modeling and Targeting Pancreatic Cancer, Munich, Germany; Else Kröner Clinician Scientist Professor for Translational Pancreatic Surgery, Munich, Germany

**Author notes:** Corresponding author: O. Ugur Sezerman; Department of Biostatistics and Medical Informatics, School of Medicine, Acibadem Mehmet Ali Aydinlar University, Istanbul, Turkey.

**Keywords:** Pancreatic Cancer, Molecular Subtypes, Tumor Stroma, Migration and invasion, Perineural Invasion

## Abstract

We investigated the association between pancreatic ductal adenocarcinoma (PDAC) subtypes and neural invasion (NI) status in order to assess what factors influence the inter-tumor heterogeneity of NI. We built single-sample classifiers for prominent subtype schemes using multiple approaches. We predicted the subtype labels of samples with known NI status using the classifiers based on pairwise gene expression comparisons, which were consistently the best-performing models. These classifiers were made available in a Shiny app (https://salihaey.shinyapps.io/pcsubtypes/). Stromal subtypes were found to be most significantly associated with NI, with activated stroma being enriched in neuro-invasive samples. We compared the results of differential expression and gene set enrichment analyses between NI and stromal subtypes from both bulk and single-cell RNA-seq data, which revealed a prominent role of myCAFs and common signatures between activated stroma and NI (e.g., ECM remodeling and TGFβ signaling) as well as signatures that changed in divergent directions for activated stroma and NI (e.g., iCAF markers). Our study presents support for the clinical implications of PDAC stromal subtypes by revealing their association with NI incidence, while providing biological insights into the interactions between diverse genetic programs that influence NI emergence.

## 1. Introduction

Pancreatic ductal adenocarcinoma (PDAC) has several distinguishing clinical characteristics, including poor prognosis, early distant metastases, and resistance to conventional treatment options. Another hallmark of PDAC is neural invasion (NI), also known as perineural invasion, in which cancer cells surround at least 1/3 of the circumference of a peripheral nerve or invade any of the three connective tissue layers of the nerve sheath (epineurium, perineurium, and endoneurium) [1–4]. Among the cancer types that are known to display NI, PDAC has been reported with the highest incidence and severity rate [3, 5, 6]. NI is associated with recurrence, poor prognosis, and pain, and is also considered to be an independent path to metastasis, since it can exist as a distinct pathology and comprise the only route to tumor dissemination in the absence of lymphatic or vascular invasion [2–5, 7]. The underlying molecular mechanism of NI is an ongoing research area, as it has so far been found to be a multistep process that involves coordinated signaling between cancer cells and several other actors [2, 8–11]. Factors that determine inter-tumor heterogeneity regarding NI incidence are especially unclear, even though PDAC has well-established transcriptional molecular subtype classifications. The subtypes defined by Collisson et al. (classical, quasi-mesenchymal – QM, and exocrine-like) [12], Moffitt et al. (classical and basal-like tumor; activated and normal stroma) [13] and Bailey et al. (squamous, immunogenic, pancreatic progenitor and aberrantly differentiated exocrine – ADEX) [14] have been extensively studied and have been replicated by external efforts [15–17]. Furthermore, the independent prognostic value and complementarity of the Moffitt and Bailey classifications have been validated with multiple datasets [17].

The clinical value of PDAC subtyping as a treatment response signature has been noticed since several studies reported Moffitt’s basal-like subtype to be more chemoresistant compared to the classical subtype [18–20]. In order to explore the association of different subtypes with further clinical or cellular characteristics, it has been necessary to implement the established subtyping approaches on previously unseen samples. The most popular approach has been unsupervised clustering of new cohorts using subtype-specific gene signatures to determine the labels of new samples [21, 22]. However, as this approach is dependent on the sample distribution of the cohort and is unfeasible for small sample sizes, single-sample classifiers are also useful, such as PurIST [20] or other methods that rely on within-sample gene expression comparisons [23].

In this work, to assign subtype labels to samples with known NI status, we have trained cohort/platform-independent single-sample classifiers for Collisson, Moffitt tumor, Moffitt stroma and Bailey schemes and made the models available as a Shiny app (https://salihaey.shinyapps.io/pcsubtypes/). We found NI to be associated with Moffitt’s activated stroma subtype and validated this finding with the implementation of another subtyping scheme that also distinguishes activated stroma [24]. Finally, we explored the shared underlying mechanisms between stroma activation and NI by comparing the gene expression programs that are involved and found an interplay of complex processes that likely contribute to the relationship between tumor microenvironment and tumor cell invasiveness.

## 2. Materials and Methods

### 2.1. Data Preprocessing

For the Collisson discovery dataset, robust microarray analysis normalized expression values for 66 human PDAC samples were retrieved from Gene Expression Omnibus (GEO) (<u>GSE17891</u> and <u>GSE15471</u>). For Moffitt data, quantile normalized expression values for the 145 primary PDAC samples were used as submitted to GEO (<u>GSE71729</u>). For Bailey subtypes, 69 of the 96 samples from the discovery dataset had fragments per kilobase per million reads (FPKM) values available in the International Cancer Genome Consortium (ICGC) data portal [25] (https://dcc.icgc.org/). 25 other samples had quantile normalized log2 transformed microarray data available, making a total of 94 samples. For The Cancer Genome Atlas (TCGA; https://www.cancer.gov/tcga) data, raw gene expression counts (for differential expression analysis) and FPKM values (for training classifiers based on within-sample gene expression rank) were downloaded from the Genomic Data Commons Data Portal (https://portal.gdc.cancer.gov/). For the Renji dataset [26], 50 raw Agilent files were retrieved from GEO (<u>GSE102238</u>), and the data was imported, background corrected and quantile normalized using the limma R package [27]. Probe annotation was achieved by mapping the Agilent-052909 CBC_lncRNAmRNA_V3 microarray probe sequences onto NCBI RefSeq [28] Select RNA sequences (https://ftp.ncbi.nlm.nih.gov/blast/db/refseq_select_rna.tar.gz; retrieved on 15.12.2023) using the rBLAST R package [29] with default parameters. Hits were filtered for 100% identity and alignment length of 60. Only primary tumor samples were included for analyses. NI labels for 159 TCGA-PAAD samples were obtained from digitized pathology reports.

### 2.2. Classification

We built single sample classifiers for Moffitt tumor, Moffitt stromal, Bailey and Collisson subtyping schemes according to within-sample gene expression comparisons (“gene1 < gene2”) using the pair-based Random Forest (RF) approach implemented in the multiclassPairs R package [30]. For training, we used the data and labels from the respective publications of each scheme, and previously discovered [31] TCGA subtype labels for Collisson, Moffitt tumor and Bailey subtypes. We used a random 80-20 split of the samples of each dataset for training and validation. Before determining classification rules, we sorted the top 2500 most variable genes in each dataset according to importance (Gini impurity) both in distinguishing samples of each class from the rest, and in separating all classes from each other, which generated multiple lists. We picked the same number of top genes (50,100,150 or 200 – depending on the number of subtypes in the scheme) from each list so that the number of unique genes in the combined list is above 200, which allowed the number of binary combinations of genes for rules to be at least 19900. We then sorted the rules according to Gini impurity both by sample and overall and picked the number of rules from each list that performed the best via cross-validation. The top 30 rules from each list (without allowing gene repetition) were used to construct the final RF model, with the Boruta filtering turned on to eliminate less informative rules.

For the centroid classifier we used the following signature genes: Moffitt’s 25 top ranked genes each from the “basal-like”, “classical”, “activated stroma” and “normal stroma” NMF factors, and Collisson’s 62 and Bailey’s 613 differentially expressed genes (DEGs) from their multiclass SAM analyses.

For the pathway transformation-based classifier, we transformed single sample gene expressions to pathway enrichment scores (MSigDB database [32] v7.0 “all canonical pathways”) using gene set variation analysis [33], and trained each RF model with 5000 trees.

### 2.3. Bulk Transcriptomics Analyses

We performed differential expression analysis using the following combinations: 1) for Moffitt microarray data, between active stroma versus normal stroma samples using limma, and 2) for TCGA RNA-seq data, between neuro-invasive versus non-invasive samples using DESeq2 [34]. DEGs (P-adj < 0.05) from individual analyses and the genes that we found to be differentially expressed both between Moffitt stroma subtypes and for NI status were used for gene set enrichment analysis (GSEA). We used the MSigDB database v7.0 “all canonical pathways” gene set and the fgsea R package [35] for its implementation of pre-ranked GSEA. We used the ESTIMATE method in the immunedeconv R package [36] to estimate tumor purities for TCGA samples. All analyses were conducted using R version 4.2.0.

### 2.4. Single-cell RNA-seq Analysis of Stromal Subtypes

Single-cell RNA-seq (scRNA-seq) Seurat [37] objects for stroma, myeloid, lymphocyte, epithelium and endothelium cell subsets integrated from multiple datasets were retrieved from OH et al. [38] source data site (http://pdacr.bmi.stonybrook.edu/scRNA/). Pseudobulk gene expression values for each cell type were calculated by aggregating cell expressions for each sample (10 activated stroma, 16 normal stroma) by using Seurat’s SplitObject function. DEG analysis was performed for each cell type that had more than 3 samples for each of activated and normal stromal subtypes using DESeq2. For visualization, gene set signature scores for each cell were calculated by the mean expression of the gene set genes in the cell.

## 3. Results

### 3.1. Subtype Classification

Previous studies have established clinically and biologically relevant PDAC subtypes. In order to assign new samples to these existing schemes, we have trained single-sample classifiers using three different approaches. The first approach comprised of determining centroids for each subtype using the signature genes from each scheme by gene-wise centering of the expressions. New samples were assigned to the centroid subtype with the highest Spearman’s correlation coefficient. The second approach was based on the comparison of within-sample expressions of gene pairs and building RF models from the binary rules (see Materials and Methods). The final approach was to transform each sample’s gene expression values to pathway enrichment scores using gene set variation analysis, then training RF models using the pathway enrichment scores.

We have applied all approaches for generating classifiers for Collisson’s, Bailey’s, Moffitt’s tumor, and Moffitt’s stromal subtypes. For training, we either only used the subtype discovery data from the original publications (for centroid-based approach) or combined discovery data with TCGA PAAD data (for RF based approaches) if the subtype label for a sample was available [31]. We randomly generated 80/20 training-validation data splits and checked the concordance between previous subtype labels and our predictions. To confirm that the performance metrics were reproducible, we conducted 5 random splits per scheme (**Fig. 1A, Supplementary Table S11-S12**). The binary rule RF models resulted in mean weighted average F1 scores (95% confidence) of 0.88 ∓ 0.06 (Collisson), 0.89 ∓ 0.04 (Moffitt tumor), 0.92 ∓ 0.02 (Moffitt stromal), and 0.75 ∓ 0.06 (Bailey).

**Figure 1:**
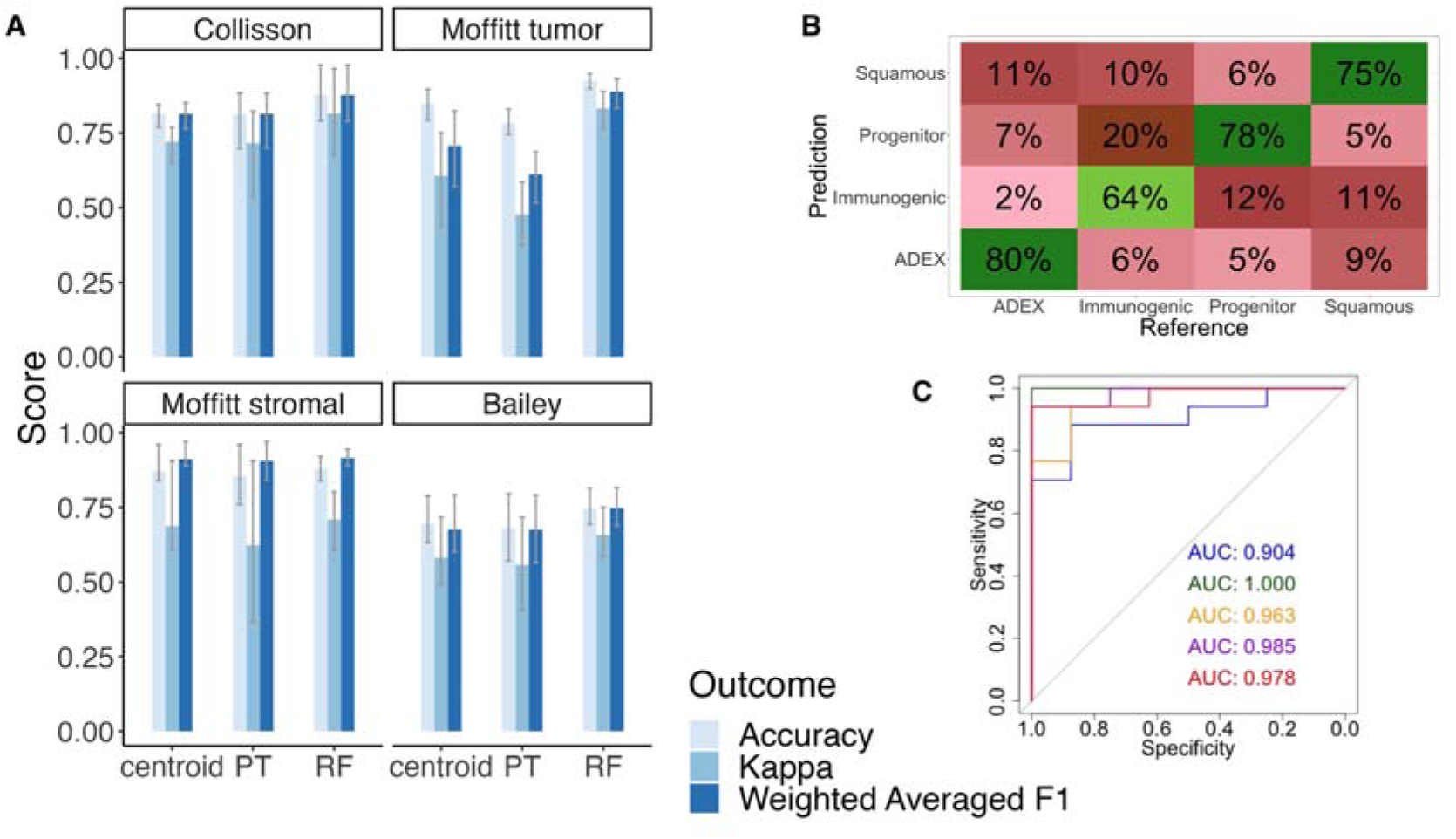
Single-sample classifiers using gene expression were trained to assign new samples to PDAC subtypes. Model performances were based on five random 80/20 splits of the training data. A) Bar plots of mean performances (accuracy, kappa, weighted averaged F1) of three different classification approaches (centroid, pathway transformation -- PT, binary rule random forest -- RF) for each subtype scheme show that binary rule RF models outperform the other models. B) Confusion matrix of the binary rule RF classifier’s test results for the Bailey scheme indicates lower accuracy for the immunogenic class. C) Receiver operating characteristic curves of the binary rule RF models with different data splits for the Moffitt stroma scheme confirm that the classifier performance is reproducible for different random data splits (AUC: Area under the ROC Curve).

The lowest performing scheme was Bailey. The binary rule RF models performed well in separating progenitor and squamous samples, but some samples of these were misclassified as the immunogenic class (**Fig. 1B**), which had low mean precision (0.69) and mean recall (0.64). The highest performing binary rule RF classifier was of Moffitt stromal subtypes (**Fig. 1C**). Unlike the other schemes, Moffitt stromal models were trained using only the Moffitt discovery dataset because TCGA data were only labeled [31] for the epithelial subtypes of Collisson, Moffitt tumor and Bailey. In order to make sure the relatively lower performances of the epithelial models were not due to the inclusion of multi-platform datasets, we also trained binary RF models for those schemes using only their discovery datasets. The discovery-only model performances were lower than the models with TCGA data included (**Supplementary Figure 6**).

We also compared our models’ performances to a previously published classifier [20], PurIST, for Moffitt tumor subtypes, which resulted in an F1 score of 0.69 for the Moffitt dataset, even though the Moffitt dataset was one of the training sets for PurIST. The predicted labels from PurIST and the labels we use for testing model performance (from GEO GSE71729) are different because PurIST used different subtype labels for training, generated via consensus clustering of the datasets included in their study. Throughout this work, we adhered to using the original subtype labels as provided in their respective discovery papers. As the binary rule RF models showed significantly improved concordances compared to the centroid and pathway transformation models, we predicted the subtypes for unlabeled samples using these models (**Fig. 2A**). We confirmed via cross-prediction and multiple correspondence analysis (**Fig. 2B**) that Bailey’s squamous and Collisson’s QM samples strongly overlap with Moffitt’s basal-like, while Bailey’s progenitor and Collisson’s classical samples strongly overlap with Moffit’s classical subtypes. Furthermore, Collisson’s exocrine-like samples strongly overlap with Bailey’s ADEX samples. We performed another associative analysis using only the Collisson, Moffitt and Bailey datasets with their original training labels alongside predicted labels from alternate schemes. In this analysis, for each sample, only the original discovery labels were known, and the labels for the other schemes were predicted by our models. Therefore, the associations are contingent on the accuracy of the models. Inspecting standardized Pearson residuals (PR) confirmed that samples that are known to belong to classical, basal-like or exocrine-like groups are assigned by our models to the corresponding subtypes in alternate schemes, as indicated by the strong associations (PR > 4) with known cross-scheme subtype correspondences (**Fig. 2C**). Additionally, we verified the independence of Moffitt’s stromal and epithelial subtypes using our predicted labels for the TCGA, Collisson and Bailey datasets (Chi-squared test P = 0.84, **Fig. 2D**), making sure that the association was dependent on the accuracy of our predicted labels.

**Figure 2:**
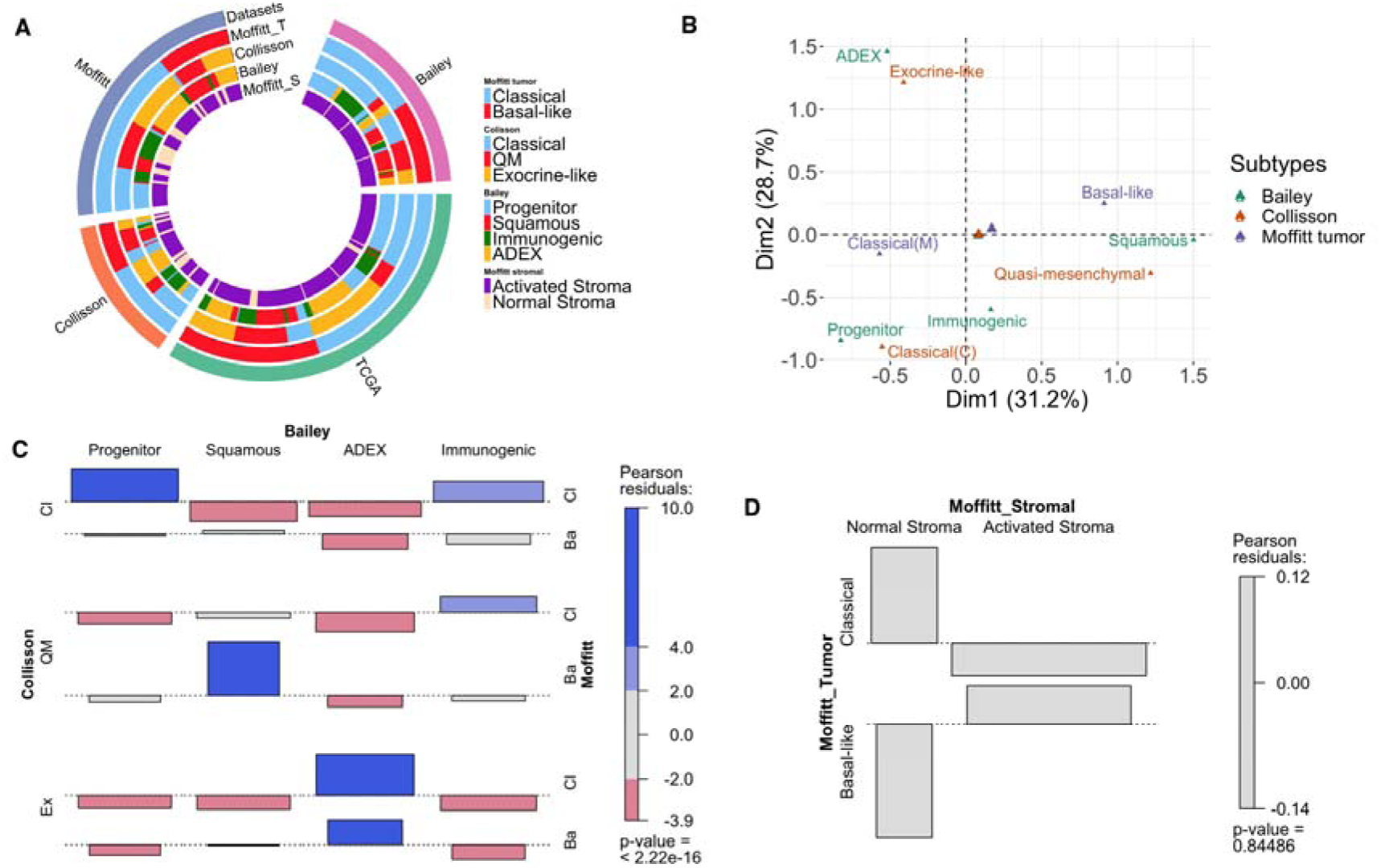
Cross-prediction of subtype discovery and TCGA datasets using single-sample subtype classifiers revealed overlaps between subtypes of different schemes. A) Circular heatmap showing subtype labels for all samples in subtype discovery and TCGA datasets. B) Multiple correspondence analysis plot of Collisson, Bailey and Moffitt epithelial subtypes using subtype discovery and TCGA datasets (n=460) shows that subtypes from different schemes that correspond to classical, basal-like, or exocrine-like characteristics are grouped together; Classical group: Collisson classical, Moffitt classical, Bailey pancreatic progenitor; Basal-like group: Collisson quasi-mesenchymal, Moffitt basal-like, Bailey squamous; Exocrine-like group: Collisson exocrine-like, Bailey ADEX. C) Standardized Pearson residuals from associative analysis for epithelial subtypes using Collisson, Moffitt and Bailey discovery datasets (n=283) confirm that the predicted labels for the samples that belong to classical, basal-like, or exocrine-like subtypes are in corresponding epithelial groups to the discovery labels. (Ba: basal-like, Cl: classical, QM: quasi-mesenchymal, Ex: exocrine-like) (Chi-squared test P < 2.26e-16) D) Associative analysis for Moffitt tumor and stromal subtypes using TCGA, Collisson and Bailey datasets (n=337) confirms the independence of Moffitt’s stromal and tumor subtypes (Chi-squared test P = 0.84).

We also performed one-way ANOVA to assess the differences in tumor purities across subtypes using TCGA data, confirming [31] that Moffitt tumor classification was independent of purity (*p-value* = 0.26) and Moffitt stroma classification was also independent of tumor purity (*p-value* = 0.10). Significant differences in purities between groups were observed for both Collisson (*p-value* = 1.49e-05) and Bailey (*p-value* = 9.94e-11) subtypes. For Collisson subtypes, post-hoc Tukey’s Honest Significant Difference (TukeyHSD) test revealed the classical group had significantly higher mean purity than the exocrine (*p-value* = 1.07e-04; diff = -0.12) and QM (*p-value* = 1.20e-4; diff = -0.14) groups. For Bailey subtypes, TukeyHSD analysis showed that the progenitor group exhibited significantly higher mean purity compared to the ADEX (*p-value* = 6.23e-06; diff = -0.14) and immunogenic (*p-value* = 7.72e-10; diff = -0.22) groups, and the squamous group also had significantly higher mean purity than the ADEX (*p-value* = 0.014; diff = -0.11) and immunogenic (*p-value* = 1.83e-5; diff = -0.18) groups, with no significant difference observed between the squamous and progenitor groups (*p-value* = 0.61). Lastly, one-way ANOVA revealed that NI status was independent of purity (p-value = 0.5).

### 3.2. Activated Stroma is Associated with NI

Next, we predicted missing subtype labels in the TCGA dataset using our models and explored the impact of molecular subtypes on NI status. Among the 159 samples with NI status included in the TCGA digitized pathology reports, 25 were indicated to display no NI, while the rest were found to be neuro-invasive. We also predicted subtypes for the samples in the Renji cohort for which NI status and gene expression microarray data is available in GEO (GSE102238). Within the 50 samples of the Renji cohort, 22 were classified as non-invasive, and 28 were labeled as neuro-invasive. Among the four subtyping schemes we considered (**Fig. 3A-D**), the classification scheme that was most strongly associated with NI status was Moffitt stromal (n = 209; Fisher’s exact test *P =* 1.93e-05; **Fig. 3D**). Inspecting PRs using association plots [39] revealed that non-invasive samples’ concurrence with the normal stroma subtype significantly (PR > ±2) deviates from the expected value (PR *=* 3.52; **Fig. 3E**). A Cochran-Mantel-Haenszel test was used to check that the observed association was consistent across both datasets, which yielded a p-value of 0.047. Even though the result was significant, we also checked the associations within TCGA and Renji datasets individually. The TCGA dataset by itself also showed that the NI status was most strongly associated with Moffitt stromal subtypes (n = 159; Fisher’s exact test *P =* 0.015; **Supplementary Fig. 1**), and that the non-invasive samples’ correspondence with the normal stroma subtype significantly deviates from the expected value (PR *=* 2.49; **Supplementary Fig. 2**). The Renji cohort by itself was also most strongly associated with Moffitt stromal subtypes, but the association was not significant (n = 50; Fisher’s exact test *P =* 0.537; **Supplementary Fig. 3**). The PRs for this cohort were similarly not significant, but changed in the same direction as TCGA findings, i.e. non-invasiveness positively associated with normal stroma and neuro-invasiveness with activated stroma (**Supplementary Fig. 4**). Therefore, as the associations from the individual datasets maintained the same directionality, concatenation of the labels of the two cohorts had consolidated the findings from the individual datasets.

**Figure 3:**
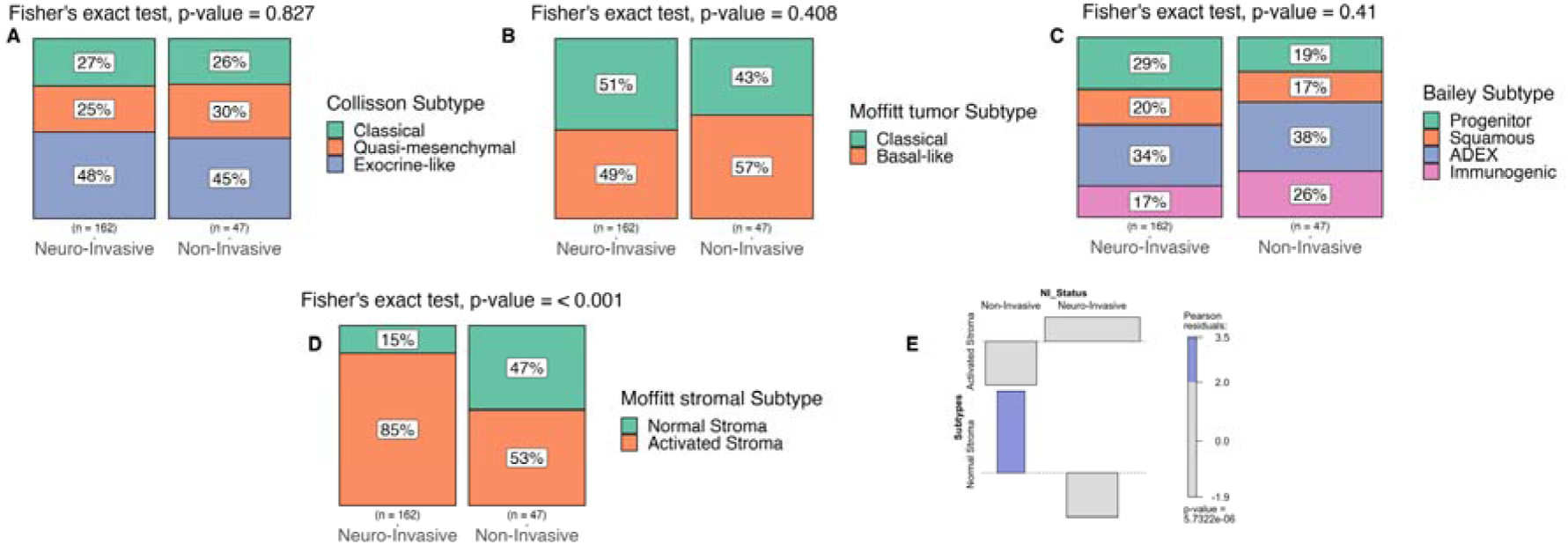
A-D) Mosaic plots showing NI status distribution among subtypes of the TCGA and the Renji datasets, and the corresponding Fisher’s exact test results (n = 209). The classification scheme that was most strongly associated with NI is Moffitt stromal (D). E) Standardized Pearson residuals (PR) from associative analysis between Moffitt stromal subtypes and NI status show that non-invasive samples are enriched in the normal stroma subtype (n = 209, PR = 3.52, Chi-squared test P = 5.73e-06).

In order to validate the association of activated stroma with NI, we also considered the subtype classifications by Puleo et al. [24]. Instead of designating orthogonal stromal and epithelial classifications, Puleo subtypes comprise a whole-tumor classification scheme that distinguishes a “stroma activated” subgroup from four others (“desmoplastic”, “pure basal-like”, “pure classical”, and “immune classical”). We assigned Puleo subtype labels to samples by using the centroid classifier provided with the original paper. Each sample’s Spearman rank correlation to the five centroids was calculated, and the centroid with the most correlation was assigned as the predicted label. As a result, the majority of samples identified as stroma-activated belonged to the neuro-invasive group. However, Fisher’s exact test across all five subtypes revealed a modest statistically significant association (n = 209; Fisher’s exact test P = 0.048; **Fig. 4A**). Further analysis focusing on the 101 samples that belonged to the three Puleo subtypes which are characterized by strong stromal signals (stroma activated, desmoplastic or immune classical), demonstrated a significant correlation with NI status (n = 101; Fisher’s exact test P = 0.01; **Fig. 4B**).

**Figure 4:**
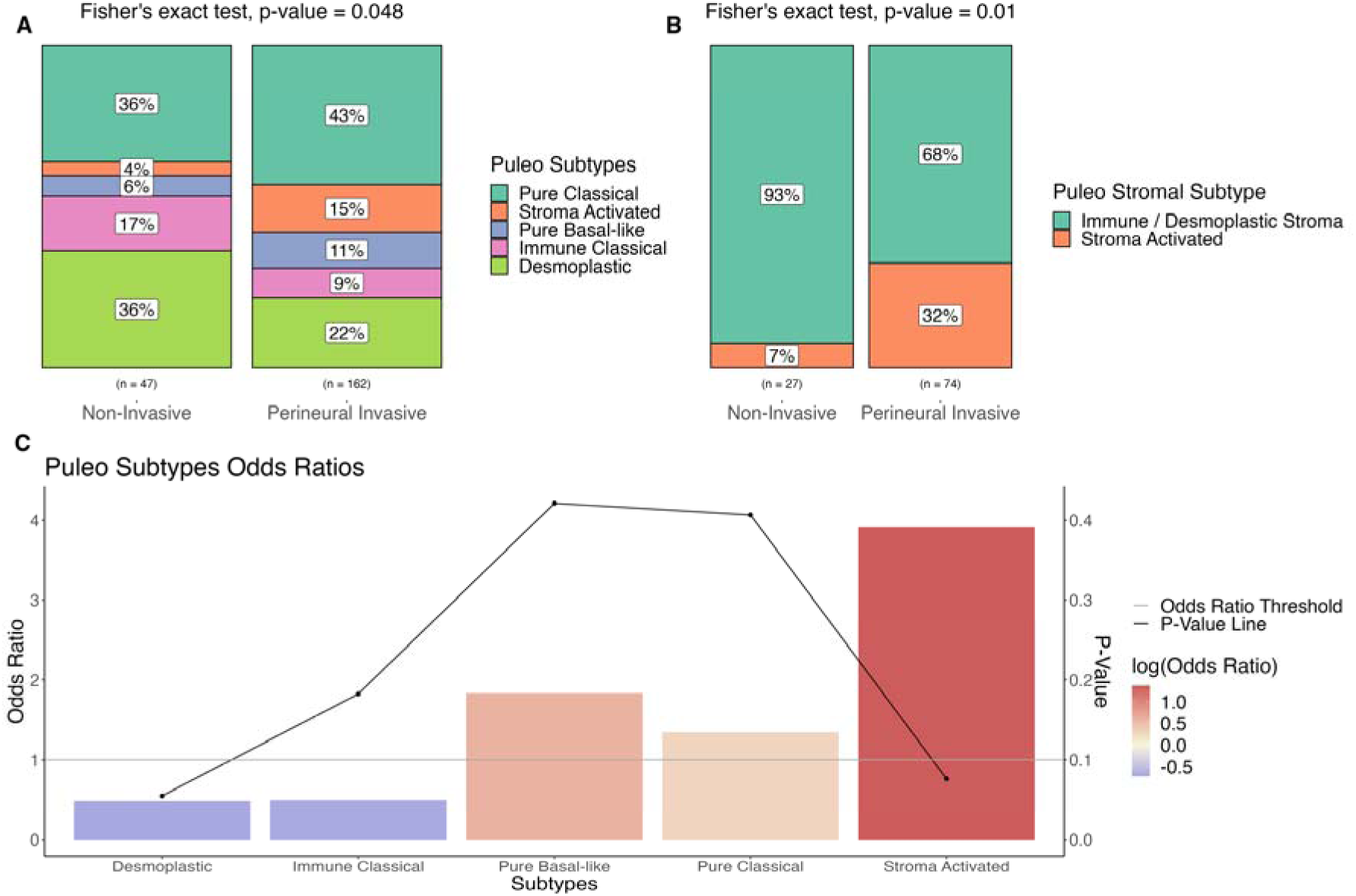
Puleo subtypes validated the association of NI with stroma activation. A) Mosaic plot of NI status distribution among Puleo subtypes of the TCGA and the Renji datasets shows that the majority of stroma activated samples are neuro-invasive (n=209). B) Mosaic plot of NI status distribution among samples that belong to Puleo subtypes that are characterized by stroma signals (stroma activated, desmoplastic or immune classical) demonstrate association with NI status (n = 101). C) Odds ratios for Puleo subtypes (> 1 indicating a higher prevalence in the NI group compared to the non-invasive group) and one-vs-rest Fisher’s exact tests (P-value) illuminate the directionalities of the associations in panel A. Stroma activated subtype is seen in neuro-invasive samples and desmoplastic and immune classical subtypes are seen in non-invasive samples.

In order to evaluate the correspondence of individual Puleo subtypes with NI, we also calculated odds ratios for each of the five subtypes (odds ratio > 1 suggesting a higher prevalence in the NI group compared to the non-Invasive group) and performed one-vs-rest Fisher’s exact tests for each subtype (**Fig. 4C**). This showed that the stroma activated subtype, which corresponds to the Moffitt’s activated stroma, mostly occurred in neuro-invasive samples (Odds Ratio = 3.91, Fisher’s exact test P = 0.076), while the desmoplastic subtype, corresponding to the normal stromal subtype, was mostly seen in non-invasive samples (Odds Ratio = 0.49, Fisher’s exact test P = 0.055).

### 3.3. Comparison of Gene Expression Programs Between Stroma Activation and NI

To characterize the shared genetic programs between stroma activation and NI, we performed DEG and GSEA analyses between active stroma versus normal stroma samples and between neuro-invasive versus non-invasive samples (**Fig. 5; Supplementary Table S1-S5**). Among the 4673 stroma and 2807 NI DEGs that were found to be significant (P-adj < 0.05), 335 were found to be common and changed in the same direction. Furthermore, between 125 stroma and 216 NI significantly enriched pathways (P-adj < 0.05), 20 were common and changed in the same direction. Among the enriched signatures from common DEGs, the top ones were matrix metalloproteinases, with interstitial collagenase (*MMP1*) and stromelysin-3 (*MMP11*) both upregulated. Other matrix metalloproteinases (*MMP7*, *MMP8*, etc.) were found to be upregulated in individual analysis results as well. Collagen degradation and ECM regulators pathways were also enriched independently in both analyses due to upregulation of different genes. Conversely, normal stroma and non-invasive samples were enriched in muscle contraction markers (e.g., *DES* and *VIM*). Normal stroma also showed a significant upregulation in myofibroblastic cancer associated fibroblast (myCAF) marker expressions (e.g. *ACTA2*, *TAGLN, MYL9*, *CALD1*, *TPM2*), and some inflammatory CAF (iCAF) markers such as *CXCL12*, *IL6*, *CLEC3B*, and *CCL2*. We also checked the entire set of iCAF/myCAF marker genes [40] to formally analyze iCAF and myCAF activity in stromal subtypes and NI (**Supplementary Fig. 5**). We performed GSEA for iCAF and myCAF gene sets similarly to pathway gene sets and discovered that the contrasting strong upregulation and downregulation results from different myCAF genes in stromal subtypes resulted in a non-significant result for enrichment (*p-value* = 0.18), while for neuro-invasiveness, myCAF markers were overall upregulated (*p-value* = 0.036). iCAF genes were significantly upregulated in neuro-invasive samples (*p-value* = 4.16e-4), and significantly downregulated for activated stroma samples (*p-value* = 3.92e-23). Other common signatures between the analyses included keratinization and the formation of the cornified envelope due to the upregulation of mostly the same genes. Furthermore, validated transcriptional targets of TAp63 isoforms, p53 downstream, programmed cell death, and tight junction interactions were among the prominent signals that emerged from both analyses. Common upregulation of *SPP1*, *FGF19* and *PLAU* resulted in the enrichment of the FGF pathway. There were also different genes from individual analyses that belonged to related processes, e.g., epithelial to mesenchymal transition (*CRB3*, *DSP*, *CDH1*, *WNT5A*, *SMAD3*, *TGFB2*, etc.). Finally, there were also biological process markers that changed in different directions for NI and stroma activation. For example, neuro-invasive samples showed upregulation of the cytokine-cytokine receptor interaction pathway, while activated stroma samples showed downregulation of the same pathway.

**Figure 5:**
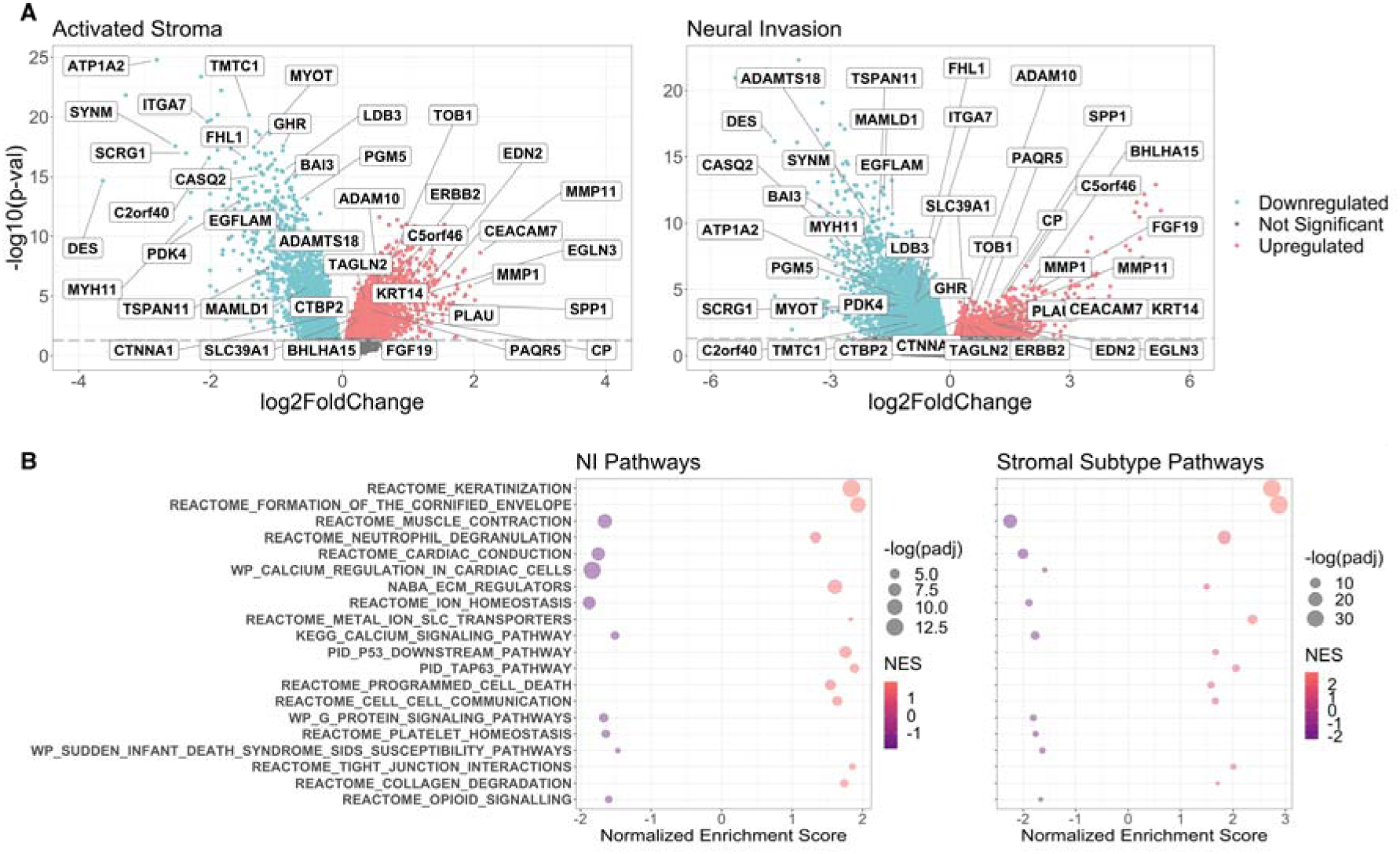
DEG and GSEA analyses for stromal subtypes (with Moffitt dataset) and NI (with TCGA dataset) demonstrated shared genetic programs between the activated stroma and NI. A) Volcano plot of genes upregulated and downregulated in stroma activation and NI; top 40 shared genes between the analyses that changed in the same direction are highlighted. B) GSEA results for pathways that are significantly enriched in both stroma activation and NI and that changed in the same direction are shown.

We also validated the clinical significance (log-rank p-value < 0.05) of 75 of the 335 common DEGs between NI and stromal subtypes by investigating the Human Protein Atlas database [41] (v23.0; proteinatlas.org) for patient survival and mRNA correlation in pancreatic cancer (**Supplementary Table S13**). We filtered the DEGs which were downregulated in NI and activated stroma by the “prognostic - favorable” statistic and upregulated DEGs by “prognostic - unfavorable” statistic.

### 3.4. Microenvironmental Influences on NI and Stromal Subtype Dynamics

In order to further explore the functional relationship between stromal subtypes and NI, we used scRNA-seq data from samples classified into Moffitt’s activated or normal stromal subtypes [38]. We investigated DEGs (P-adj < 0.05) in subpopulations of cells from stroma, myeloid, lymphocyte, epithelium, and endothelial groups (**Fig. 6; Supplementary Table S6-S10**). We also compared the bulk RNA-seq NI DEGs with the scRNA-seq stromal subtype DEGs to predict the likely cell type sources of the mechanisms inferred from bulk RNA-seq. Among the 105 total (58 unique) common DEGs that changed in the same direction for NI and activated stroma, the top cell types that represented the DEGs were of stromal origin; myCAFs showed 18.1% of DEGs, smooth muscle pancreatic stellate cells (smPSCs) showed 17.1%, quiescent pancreatic stellate cells (qPSCs) represented 16.2%, and complement-secreting cancer associated fibroblasts (csCAFs) showed 12.4% of DEGs. Myocyte, Schwann and iCAFs cells clusters were not observed in all patients due to their small population sizes and therefore did not display many significant DEGs. Neoplastic cells were also representative of a high proportion of DEGs, with 10.5%.

**Figure 6:**
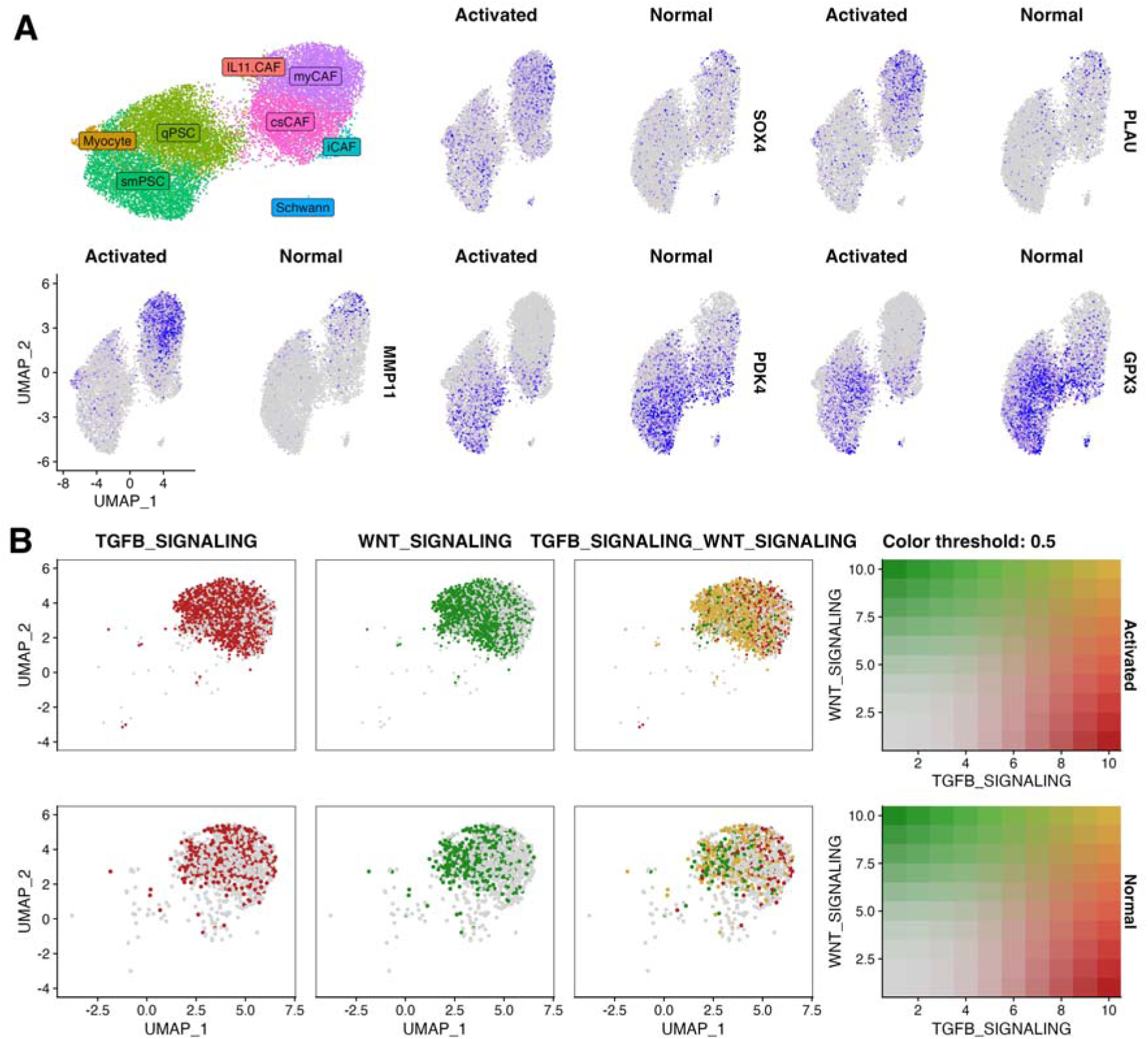
Single-cell RNA-seq analysis of DEGs between activated and normal stroma samples across major cell types highlights that stroma cells, particularly myCAFs, display signaling patterns observed in bulk analyses. A) Feature plots with uniform manifold approximation and projection for the stromal cell subset are shown, with the top left plot indicating clusters for stromal cell types; for each feature, gene expressions are displayed separately in cells from activated stroma samples and normal stroma samples for genes that were seen to be differentially expressed across multiple cell types. B) Signal scores for TGFB signaling and WNT signaling are presented both separately and in aggregate for myCAF cells from activated stroma samples (top) and in myCAF cells from normal stroma samples (bottom).

*SOX4*, *PLAU* and *MMP11* were upregulated in multiple cell types amongst different major cell groups (**Fig. 6A**). There were also cell-type specific DEGs, including upregulation of *SPP1* in classically activated monocytes, downregulation of *DMPK* (involved in muscle contraction) in smPSCs, or downregulation of *NPDC1* (involved in neural proliferation) in qPSCs. *ITGA7* was downregulated in all major stroma cell subpopulations (myCAF, smPSC, qPSC, csCAFs), while *ITGAV* was only upregulated in qPSCs. Neoplastic cell DEGs that were shared with NI were mostly cell-type specific, including upregulation of *CALB2* and downregulation of *PCP4* (involved in neuron differentiation). Some pathways emerged as common between NI and activated stroma due to upregulation of different genes (**Fig. 6B**); TGFB signaling and SMAD2/SMAD3 pathways in myCAFs via upregulation of genes including *INHBA*, *TGFBR1*, and *SMAD2*, and in NI through upregulation of *TGFB2*, *MMP1*, *ATF3* and *SMAD3*. Early (pre-activation) CD8+ T-cells showed upregulation of Activator Protein-1 (AP1) family member *FOSB*, and NI showed upregulation of *JUN*, another AP1 protein, both involved in TGFB processes. Similarly, besides *PLAU* (upregulated in smPSCs, csCAFs, lymphocytes, and epithelial cells) and *ITGAV*, there were other upregulated genes involved in VEGF-A/VEGFR2 signaling both in NI (e.g. *ADAM10* and *IQGAP1*), as well as various genes in myCAFs (e.g. *P4HB*, *FSCN1* and *CALU*), smPSCs (e.g. *VEGFA*), and in csCAFs (e.g. *SSR3*, *ITGB5* and *MMP14*). Furthermore, Besides *SOX4* upregulation in many cell types (all major stromal cell types, lymphocytes, tumor associated macrophages, and endothelial cells), various genes involved in WNT signaling pathway (*WNT2*, *LEF1*, or *NKD2*) were upregulated in csCAFs. There were also new DEGs involved in programs that were already found to be common in the bulk analyses, such as *SPARC* in myCAFs, *LUM* in smPSC, *FN1* in qPSC, all involved in ECM organization.

## 4. Discussion

In this work, we found an association between NI and PDAC stromal subtypes according to two independent subtype apportionments. Our analyses revealed that non-neuro-invasive samples were enriched in Moffitt’s normal stroma subtype, and Puleo’s immune classical and desmoplastic subtypes, which were found to be corresponding to Moffitt’s normal. [24] The desmoplastic reaction, a hallmark of PDAC, may cause the cancer cells to become embedded in a dense, fibrous stroma, which may restrict the movement of the cells and make the stroma less hospitable for NI. Although desmoplasia is known to be a barrier to therapeutics, removal of stromal cells [42] that cause it or antagonizing it via the inhibition of Sonic hedgehog signaling [43] results in more aggressive tumors and decrease in survival. On the other hand, we found activated stroma to be associated with neuro-invasive samples, possibly due to effectors such as growth factors or proteases, which may support cancer cell movement and invasion.

When we compared gene expression differences to understand the mechanism behind NI and activated stroma association, we found signals that point to a complex and dynamic process. For example, in both analyses, there was an abundance of genes related to extracellular matrix organization and collagen degradation, which may be showing that stromal changes facilitate cancer cell movement and NI through ECM remodeling. Aberrant activation of growth factors like the FGF pathway may also be promoting NI through crosstalk with various other pathways, such as TGFβ, which was upregulated in neuro-invasive samples. Furthermore, upregulation of TGFβ signaling pathway genes also featured prominently in the stromal subtype scRNA-seq analysis. Alongside TGFβ signaling, SMAD2/SMAD3 pathways, the VEGF/VEGFR2 signaling axis, and WNT/β-catenin signaling were identified as other common mechanisms between activated stroma and NI, with a diverse array of cell types within the activated stromal microenvironment showing upregulation in different genes. These parallel changes suggest the possibility of significant crosstalk between these pathways in the tumor microenvironment which may play a role in the regulation and propagation of NI.

Even though our findings are based on high-throughput data analysis and do not involve targeted validation, individual genes and pathways we discuss collectively in the context of activated stroma have been experimentally validated to be involved in NI by other works. For example, immunohistochemistry analysis in human PDAC samples has shown that high levels of TGFβ signaling activation was positively correlated with NI [44]. In another study, reverse transcription-quantitative PCR and western blot analyses in pancreatic cancer cell lines revealed that *FGF2* levels were significantly higher in neuro-invasive groups [45]. Additionally, extensive experimental validation of *MMP-1*’s involvement in NI was performed, including *MMP-1* silencing, which repressed the migration of neuro-invasive cell line cells in a Transwell assay [46]. While these studies focused on the involvement of the relative genes in NI through individual mechanisms, such as FGF’s angiogenic effects and its function in the central nervous system, examining them together within the context of stromal activation suggests cooperating mechanisms in tumor progression and invasion through interactions.

However, stroma’s involvement in cancer cell movement and NI did not appear straightforward. The upregulation of iCAF markers or inflammatory mediators such as various cytokines were shared by NI and normal stroma, even though NI was found to be unfavorable in normal stroma. Presence of shared factors may be related to why some samples display NI in the absence of activated stroma and show that there is not a single mechanism through which the stoma orchestrates NI. When we scrutinized changes in specific CAF populations using scRNA-seq, we found that, iCAF populations were scant in the scRNA-seq dataset and the signals detected in the bulk analyses could also be due to alternative biological programs. In contrast, GSEA analysis of myCAF markers in the bulk stromal subtype dataset was inconclusive due to mixed signals of both strongly positive and strongly negative activity from different marker genes. However, comparison of scRNA-seq data between activated and normal stroma samples showed the activated stroma samples to have a higher percentage of myCAFs [38]. scRNA-seq DEG analysis also revealed myCAFs to display the most significant activity in terms of common mechanisms between stromal activation and NI, which was not apparent with bulk-only analyses.

Upregulation of p53 downstream pathway, programmed cell death or validated transcriptional targets of TAp63 isoforms in both neuro-invasive and activated stroma samples shows that these samples display worse prognosis despite containing tumor suppressing signals, meaning these tumor suppressing pathways are not always sufficient to prevent tumor growth and metastasis in the presence of other processes. Previously, ΔNp63, which is another isoform of p63 and an inhibitor of TAp63 and p53 has been found to activate the basal subtype in PDAC [47]. We also observed downregulation of neuron specific genes in activated stroma which might be connected to the neural dedifferentiation processes that have previously been implicated in NI [48]. In these processes, Schwann cells were discovered as major actors, but we could not confirm their contribution to activated stroma in our analysis because they had small population sizes within the dataset. The complex interplay between various molecular and cellular factors that contribute to tumor progression in the microenvironment highlight the need for further investigation and understanding of the specific mechanisms involved in the poorer survival of the patients.

Our analyses did not reveal any significant association between epithelial subtypes and NI. It should be noted however, that Bailey subtypes’ observed lack of significant association with NI status is based on classifications from a predictive model with lower performance. As the other two epithelial subtype schemes, Moffitt tumor and Collisson, also did not reveal a significant association with NI, we did not pursue further analysis on Bailey subtypes within the scope of this work. Even though substantial research on the clinical importance of epithelial subtypes have been produced, stromal subtypes have not gotten as much attention. The Moffitt study that established the stromal subtypes suggested that activated stroma may play a role in tumor colonization in foreign sites, as patients with activated stroma had significantly higher graft success [13]. The study also found that activated stroma was associated with worse prognosis, which is a common feature with NI. Our study implicates that further molecular mechanisms are involved in NI, while also constituting support towards stroma’s involvement in NI emergence. As NI is considered an independent path to metastasis, it is of interest to determine whether the neuro-invasiveness of tumor cells is related to alternative mechanisms beyond well-studied pathways.

In conclusion, our findings highlight the need for more research into stromal subtypes and their role in tumor progression while also providing insights into the complex interplay between molecular pathways and cellular processes that are involved in NI emergence. Understanding the molecular bases of stroma activation and NI could lead to the development of novel therapeutic approaches to improve outcomes for PDAC patients.

## Abbreviations

ADEX: Aberrantly differentiated exocrine
AP1: Activator Protein-1
csCAFs: Complement-secreting cancer associated fibroblasts
DEG: Differentially expressed gene
FPKM: Fragments per kilobase per million reads
GEO: Gene Expression Omnibus
GSEA: Gene set enrichment analysis
iCAF: Inflammatory cancer associated fibroblast
ICGC: International Cancer Genome Consortium
myCAF: Myofibroblastic cancer associated fibroblast
NI: Neural invasion
PDAC: Pancreatic ductal adenocarcinoma
PR: Standardized Pearson residuals
QM: Quasi-mesenchymal
qPSCs: Quiescent pancreatic stellate cells
RF: Random forest
scRNA-seq: Single-cell RNA-seq
smPSCs: Smooth muscle pancreatic stellate cells
TCGA: The Cancer Genome Atlas
TukeyHSD: Tukey’s Honest Significant Difference.

## Acknowledgments

S.E.Y. received scholarship from The Scientific and Technological Research Council of Türkiye (TÜBİTAK) under the Grant no. 118C262.

## Author contribution

**S.E.Y.:** Conceptualization, Data curation, Software, Formal analysis, Investigation, Visualization, Methodology, Writing-original draft. **I.E.D.:** Conceptualization, Supervision, Funding acquisition, Validation, Investigation, Writing-review and editing. **E.A:** Project administration. **D.K.:** Project administration. **O.U.S.:** Conceptualization, Resources, Supervision, Funding acquisition, Investigation, Methodology, Writing-review and editing. **G.O.C.:** Conceptualization, Supervision, Funding acquisition, Validation, Investigation, Writing-review and editing.

## Data Availability Statement

The data that support the findings of this study were derived from the following resources available in the public domain:

GEO GSE17891 (https://www.ncbi.nlm.nih.gov/geo/query/acc.cgi?acc=GSE17891), GSE15471 (https://www.ncbi.nlm.nih.gov/geo/query/acc.cgi?acc=gse15471), GSE71729 (https://www.ncbi.nlm.nih.gov/geo/query/acc.cgi?acc=GSE71729), GSE102238 (https://www.ncbi.nlm.nih.gov/geo/query/acc.cgi?acc=GSE102238), ICGC data portal (https://dcc.icgc.org/), Genomic Data Commons Data Portal (https://portal.gdc.cancer.gov/), Oh et al. scRNA-seq cell subset R objects (stroma: http://pdacr.bmi.stonybrook.edu/scRNA/Stroma_Subset2021.rds, myeloid: http://pdacr.bmi.stonybrook.edu/scRNA/Myeloid_Subset2021.rds, lymphocyte: http://pdacr.bmi.stonybrook.edu/scRNA/Lymphocytes_Subset2021.rds, epithelium: http://pdacr.bmi.stonybrook.edu/scRNA/Epi_Subset2021.rds, endothelium: http://pdacr.bmi.stonybrook.edu/scRNA/Endo_Subset2021.rds).

## Supplementary Figure Legends

**Supplementary Fig. 1:** Mosaic plots showing NI status distribution among subtypes and the corresponding Fisher’s exact test results for TCGA Data.

**Supplementary Fig. 2:** Standardized Pearson residuals from associative analysis between Moffitt stromal subtypes and NI status in TCGA data.

**Supplementary Fig. 3:** Mosaic plots showing NI status distribution among subtypes and the corresponding Fisher’s exact test results for the Renji cohort.

**Supplementary Fig. 4:** Standardized Pearson residuals from associative analysis between Moffitt stromal subtypes and NI status for the Renji cohort.

**Supplementary Fig. 5:** Differential expression analysis results for iCAF/myCAF marker genes for stromal subtypes and NI status. iCAF markers were upregulated in neuro-invasive samples and downregulated in activated stroma samples. myCAF markers were upregulated for neuro-invasive samples. For activated stroma, different myCAF markers showed strong activity in both negative and positive directions.

**Supplementary Fig. 6:** The discovery-only model performances were lower than the models with TCGA data included.

