## Supplementary figures and images for "TRANSCRIPTOMIC ANALYSIS OF THE RELATIONSHIP BETWEEN PANCREATIC DUCTAL ADENOCARCINOMA STROMAL SUBTYPES AND NEURAL INVASION"

### Supplementary Fig. 1

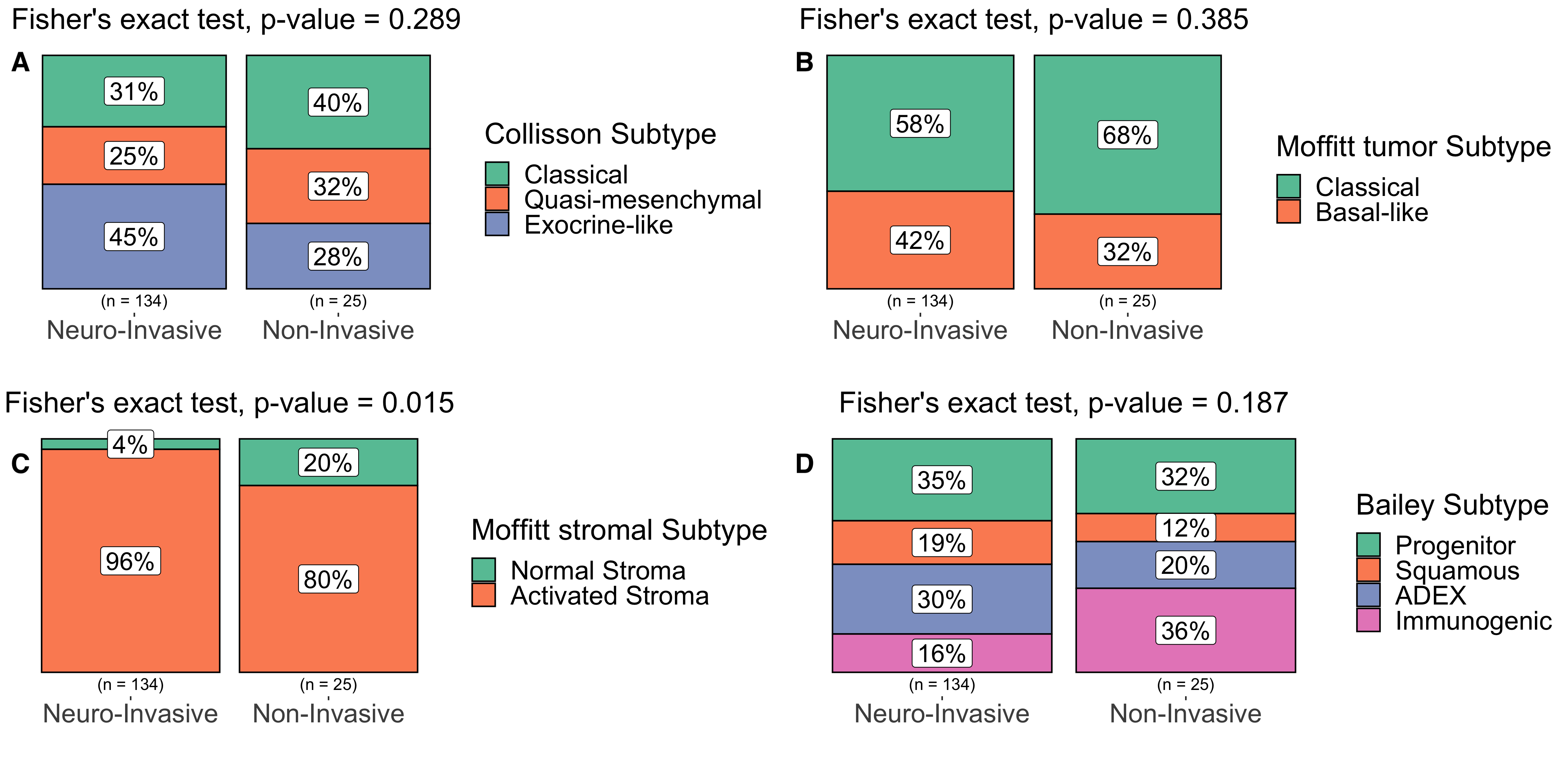

### Supplementary Fig. 2

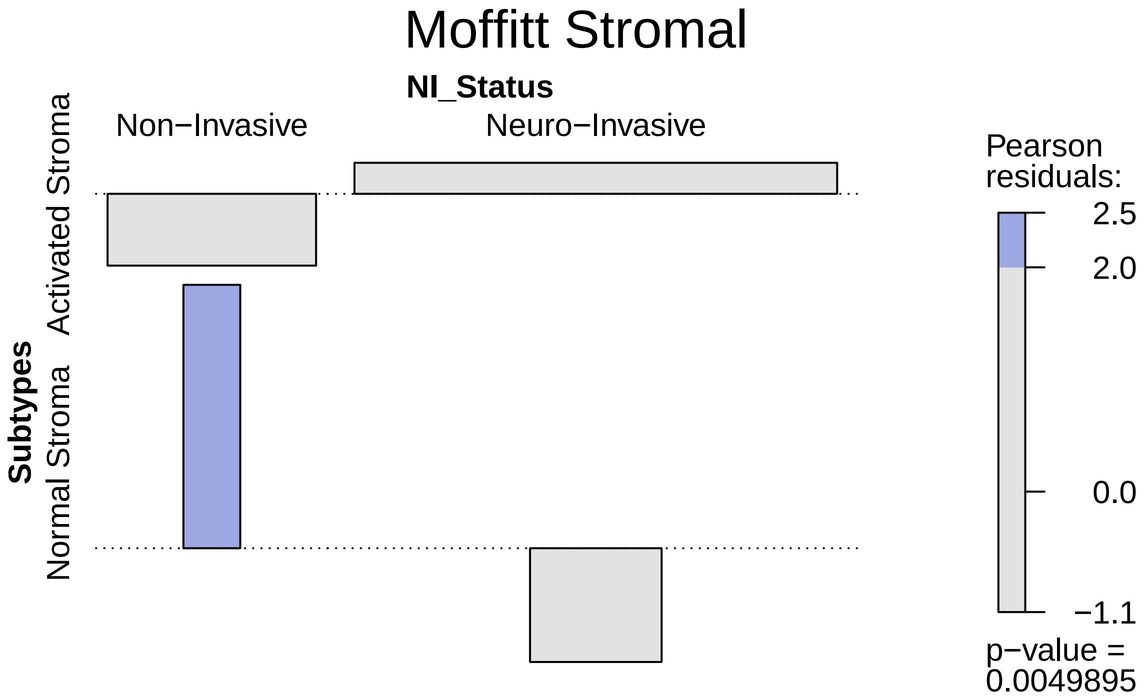

### Supplementary Fig. 3

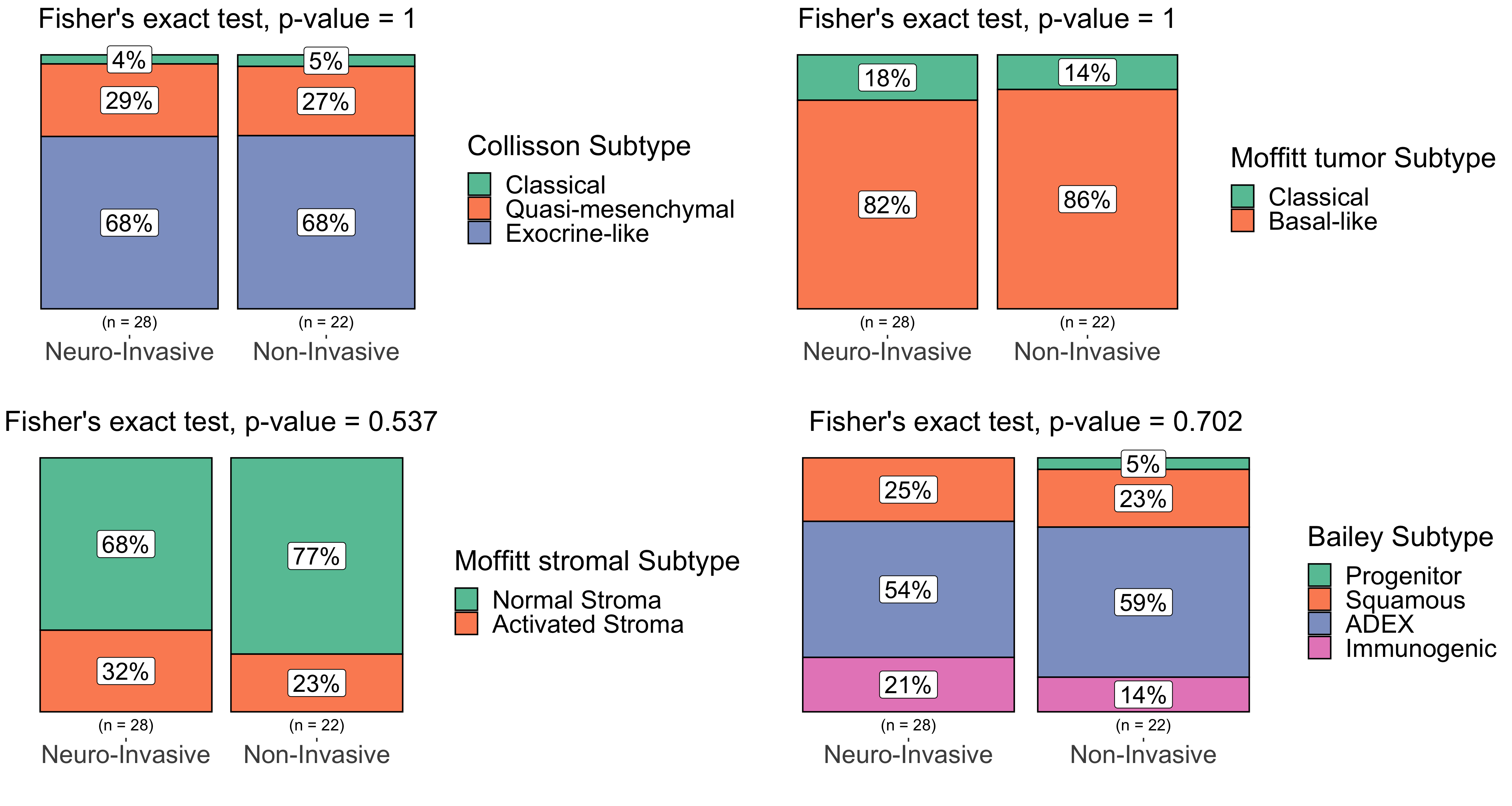

### Supplementary Fig. 4

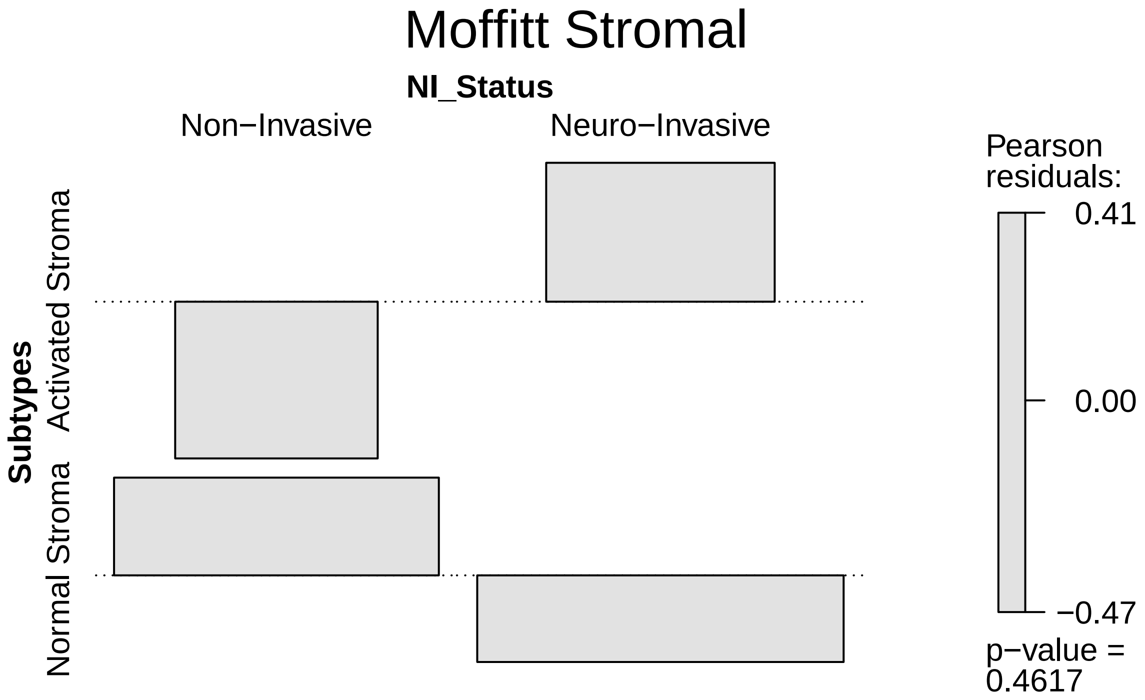

### Supplementary Fig. 5

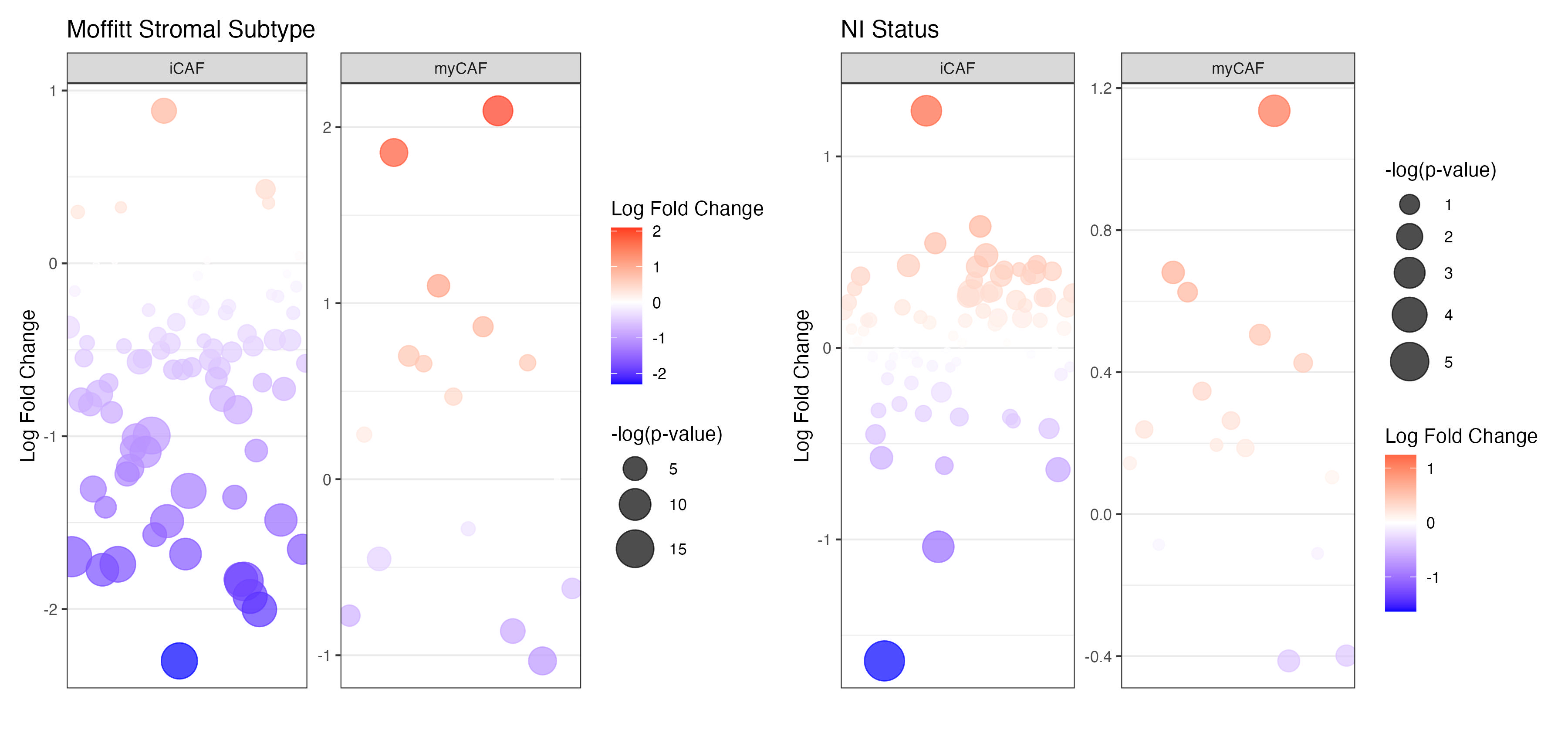

### Supplementary Fig. 6

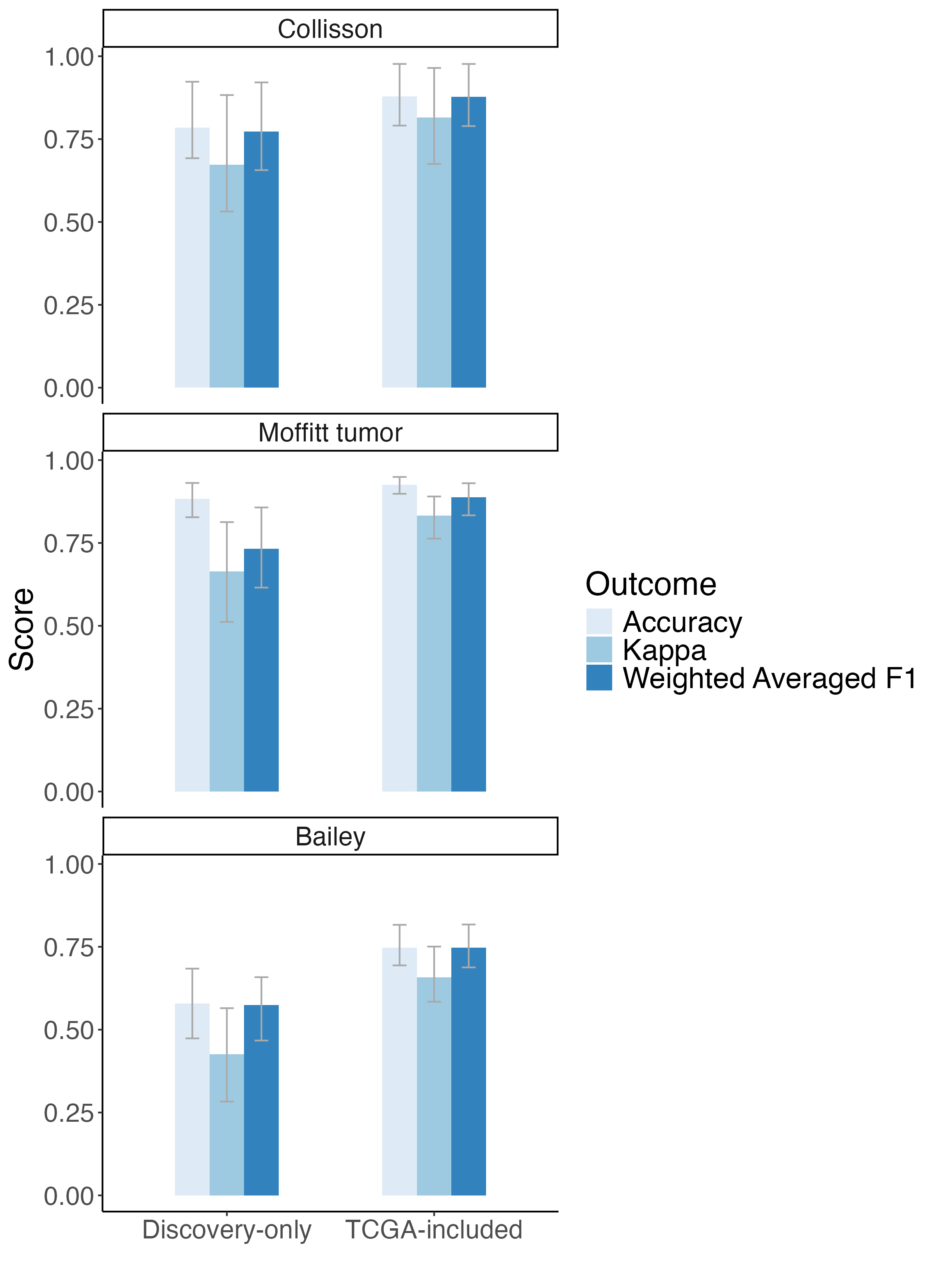
